# Novel algorithms for efficient subsequence searching and mapping in nanopore raw signals towards targeted sequencing

**DOI:** 10.1101/491456

**Authors:** Renmin Han, Sheng Wang, Xin Gao

**Author notes:** All correspondence should be addressed to Sheng Wang and Xin Gao. These authors contributed equally (Renmin Han, Sheng Wang). The program introduced in this manuscript is available at https://github.com/icthrm/cwSDTWnano.git.

## Abstract

Genome diagnostics have gradually become a prevailing routine for human healthcare. With the advances in understanding the causal genes for many human diseases, targeted sequencing provides a rapid, cost-efficient and focused option for clinical applications, such as SNP detection and haplotype classification, in a specific genomic region. Although nanopore sequencing offers a perfect tool for targeted sequencing because of its mobility, PCR-freeness, and long read properties, it poses a challenging computational problem of how to efficiently and accurately search and map genomic subsequences of interest in a pool of nanopore reads (or raw signals). Due to its relatively low sequencing accuracy, there is no reliable solution to this problem, especially at low sequencing coverage.

Here, we propose a brand new signal-based subsequence inquiry pipeline as well as two novel algorithms to tackle this problem. The proposed algorithms follow the principle of subsequence dynamic time warping and directly operate on the electrical current signals, without loss of information in base-calling. Therefore, the proposed algorithms can serve as a tool for sequence inquiry in targeted sequencing. Two novel criteria are offered for the consequent signal quality analysis and data classification. Comprehensive experiments on real-world nanopore datasets show the efficiency and effectiveness of the proposed algorithms. We further demonstrate the potential applications of the proposed algorithms in two typical tasks in nanopore-based targeted sequencing: SNP detection under low sequencing coverage, and haplotype classification under low sequencing accuracy.

## INTRODUCTION

Benefited from the deeper understanding of disease-gene associations, targeted sequencing (TS) becomes a much preferred option than whole-genome sequencing (WGS) or whole-exome sequencing because it can significantly reduce the cost, turnaround time, and data processing burden, yet provide a more focused analysis for the regions of interest typically ranging from several thousands to millions of bp. Along with the next generation sequencing, TS has been revolutionizing the way of diagnosis, prognosis, and treatment of human diseases. Oxford nanopore sequencing is a rapidly developing third generation sequencing technology that is able to generate 10-50k bp ultra-long reads in real time on a portable device at low-cost, thus provides a perfect tool for TS (Jain *et al.*, 2016; Deamer *et al.*, 2016; Stancu *et al.*, 2017). The key innovation of nanopore sequencing is the direct measurement of the electrical current signal (denoted as the *raw signal*) when a singlestrand DNA passes through the nanopore. These raw signals are transferred to reads by base-calling for further analysis.

To analyze the reads generated by nanopore-based TS, most of the bioinformatics tools follow a ‘read-to-reference’ pipeline inherited from WGS. That is, they map the base-called reads to the reference genome to locate the local genomic region of interest (Fig. 1(A)). An alternative, inverse approach is to perform subsequence inquiry of local reference genomic sequence in the ultra-long nanopore reads (Fig. 1(B)). As those reference subsequences are often known in advance with prior knowledge about the associated genes or genomic regions for the diseases, such ‘reference-to-read’ approach may overcome some challenging issues in the read-to-reference approach. For instance, in diagnostic metagenomics to detect 16S rRNA for bacteria classification, it is not necessary and very difficult to assemble the whole genomes, but TS can still detect hypervariable 16S regions in the generated reads (Fiannaca *et al.*, 2018). Also, in targeted locus amplification (TLA), a TS approach to selectively amplify and sequence entire genes on the basis of the cross linking of physically proximal sequences (De Vree *et al.*, 2014), the sequenced reads are reshuffled, therefore it is challenging for the canonical read mappers to map those reshuffled reads to the reference genome (De Vree *et al.*, 2014).

**Figure 1.**
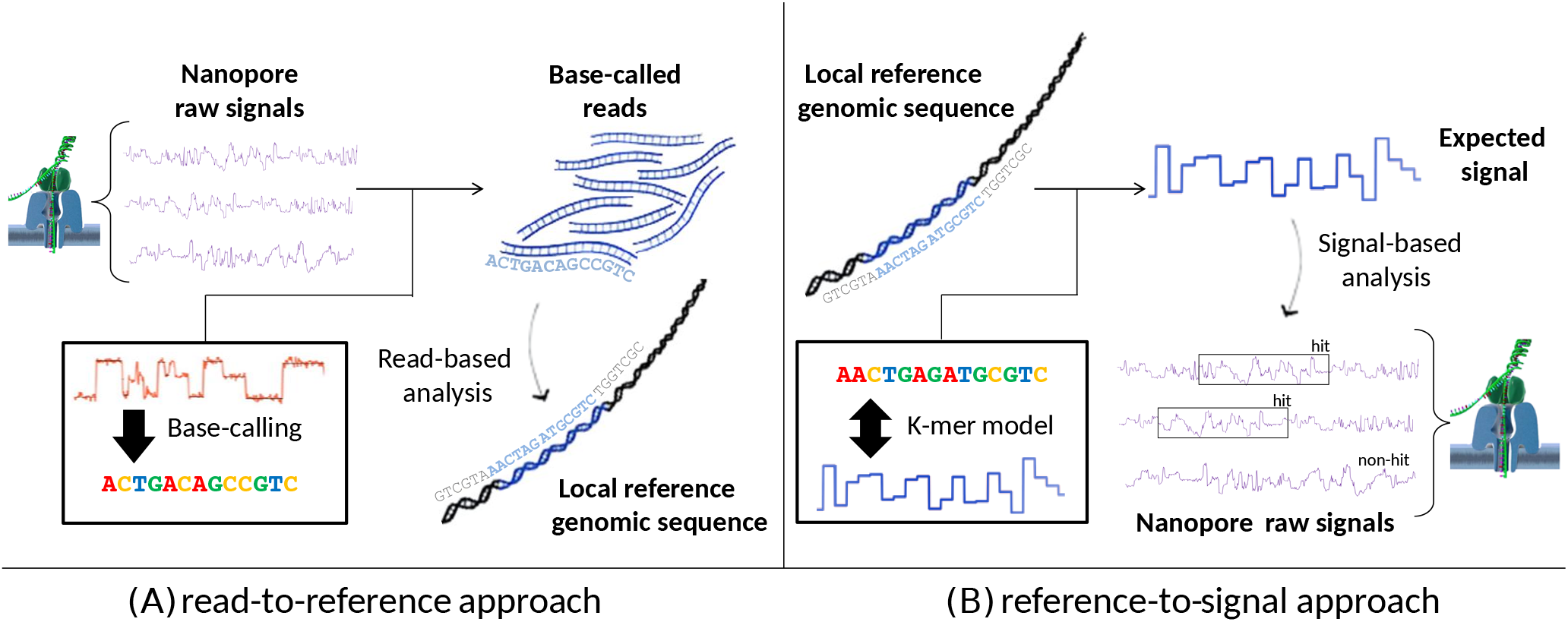
Illustration of two approaches for nanopore-based targeted sequencing. (A) The classic read-to-reference approach. In this approach, the nanopore electrical current signal sequences (i.e. raw signals) are transferred to reads by base-calling, and then the base-called reads are searched and mapped to the reference genome for further analysis (i.e., read-based analysis). (B) Our proposed reference-to-signal approach. In this approach, the local reference genomic sequence is translated to the expected signal sequence by the *k*-mer pore model, and then the expected signal sequence is searched and mapped to a pool of raw signals (i.e., signal-based analysis). The translation from the genomic sequence to the expected signals is completely reversible, while the base-calling procedure will introduce error and cause information loss in the raw signals.

Another key issue that is usually neglected in the classic read-to-reference pipeline for nanopore resides in the base-calling process, where the raw nanopore signals are translated into the nucleotide reads (i.e., ‘signal-to-nucleotide’) based on a trained machine learning model (Rang *et al.*, 2018). However, according to the recent study (Rang *et al.*, 2018), the base-calling in nanopore retains 10% to 15% error rate (Wick *et al.*, 2018), and heavily depends on the datasets that are used for training. In addition, it was found that the non-standard events, such as mutations or modifications (e.g., DNA methylation), are contained in the raw signals but lost after base-calling (Rang *et al.*, 2018). All of these defects leave a high risk of false dismissals and misalignment in local genomic region mapping. On the contrary, instead of using the base-called reads, an inverse signal-based analysis exists by first transforming the local reference genomic sequence to the expected signal sequence and then directly comparing it with the raw signals (Fig. 1(B)). The advantage of this approach resides in two folds: (i) there is no information loss in the base-calling procedure, and (ii) the transformation from the genomic sequence to the expected signal sequence is completely reversible.

In order to leverage the advantages of nanopore sequencing while avoiding its drawbacks for targeted sequencing, we propose a brand new signal-based subsequence inquiry (or reference-to-signal) pipeline that directly searches and maps a local reference genomic sequence to a pool of raw nanopore signal sequences (Fig. 1(B)). As the proposed pipeline directly operates on the raw signals but not base-called reads, and directly focus on the local region of interest, it is a more natural approach for nanopore-based targeted sequencing. There are three main benefits in this novel pipeline: (i) as the local reference genomic sequence is often known in advance, there is no need to obtain the whole reference/exon genome; (ii) because we do not perform base-calling on the raw signals, our approach has no information loss and will not miss the raw signals that contains mutations or epigenetic modifications; and (iii) the inquiry of short reference sequences will not be affected by reshuffling during TS or false dismissals caused by errors in signal-to-nucleotide translation.

However, there are several technical challenges hampering efficient reference-to-signal search: (i) the raw signal sequence is very long, often ranging from 100k to 500k bp; (ii) there is one order of magnitude scale difference between the sampling rate of the two sequences; and (iii) the alignment of real-valued sequences instead of the one of discrete letters requires accurate yet sensitive scoring functions. To our knowledge, there is no available solution to resolve these issues.

In this paper, we propose two novel algorithms to enable the direct subsequence search and exact mapping in the nanopore raw signal database (i.e., reference-to-signal). The proposed algorithms follow the principle of subsequence dynamic time warping (sDTW) and directly operate on the nanopore raw signal level. The first algorithm is the Direct Subsequence Dynamic Time Warping for nanopore raw signal search (DSDTWnano), which ensures an output of highly accurate query results and runs in an *O*(*MN*) time complexity (*M* is the query length and *N* is the raw signal length). The second algorithm is the continuous wavelet Subsequence DTW for nanopore raw signal search (cwSDTWnano), which is an accelerated version of DSDTWnano with the help of seeding and multi-scale coarsening based on continuous wavelet transform (CWT). For a typical similarity search with a 4000bp-long query and a nanopore raw signal sequence of 2105 time points, cwSDTWnano could finish the search in 600 ms. As a tool for data inquiry in targeted sequencing, two novel criteria are proposed to specify the mapping accuracy between a query genomic sequence and a raw signal sequence, which serve as the similarity measurement for the discrimination of hit and non-hit raw signals as well as the data classification.

To demonstrate the efficacy of the new approach, we make a comprehensive comparison between our reference-to-signal pipeline and the traditional reference-to-read one (using tools like BLAST (Altschul *et al.*, 1997) and minimap2 (Li, 2018)), and show that our method outperforms the traditional one by a large margin, especially when the length of the query sequence is short. We further demonstrate the potential applications of the proposed pipeline in two typical tasks in nanopore-based targeted sequencing: SNP detection under low sequencing coverage and haplotype classification under low sequencing accuracy. Results show that our algorithms achieve a very high detection and classification accuracy. Specifically, a simple SNP detection approach based on the query result of our algorithms achieves 90% detection rate under a low coverage (20×) on the E. coli dataset.

## PRELIMINARIES

### Subsequence inquiry in nanopore sequencing

As discussed in the introduction, the subsequence inquiry problem is to detect the segments of raw signals in the database that are similar to a query genomic sequence (the *hit signals*). On the contrary, the raw signals with no high-similarity segment to the query sequence are denoted as *nonhit signals*. Formally, let *X* = (*x*_1_,*x*_2_,⋯,*x_N_*) be a raw signal sequence, and *Y* = (*y*_1_,*y*_2_, ⋯,*y_M_*) be the expected query signal sequence (abbr. *query signal*) that is translated from the query genomic sequence based on the pore model (*M* < *N*). Our aim is to find a subsequence *X*[*t_s_* : *t_e_*] = (*x_t_s__*,⋯,*x_t_e__*) of *X* (1 ≤ *t_s_* ≤*t_e_* ≤*N*) that minimizes the distance measurement between *Y* and all possible subsequences of *X*:

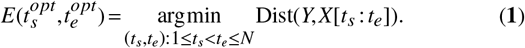

### Dynamic time warping (DTW) and subsequence DTW

Dynamic time warping (DTW) is an algorithm that measures the similarity between two temporal sequences, which is a dynamic programming technique similar to the alphabet-based alignment algorithms such as Smith-Waterman (Smith and Waterman, 1981) and Needleman-Wunsch (Needleman and Wunsch, 1970), with the distance measured by difference of the real values instead of the substitution matrix.

Given a query sequence *Y* and a database sequence *X*, the DTW distance Dist(*Y,X*) is defined iteratively:

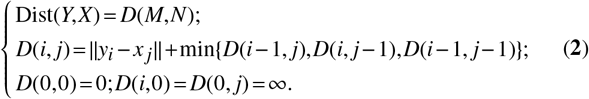

It can be seen that the DTW distance can be solved exactly in *O*(*MN*) time, resulting in the globally optimal alignment.

For the local genome-to-signal search, a naive solution is to open a sliding window for each time point in the raw signal sequence and calculate the DTW distance for each sliding window, which would result in *O*(*M*^2^*N*) time complexity, which is prohibitively high considering the large values of *M* and *N* in nanopore sequencing.

To find the optimal subsequences in an efficient way, the subsequence DTW (sDTW) (Sakurai *et al.*, 2007) is devised. By padding the query to *Y*′ = (*y*_0_, *y*_1_, *y*_2_, ⋯,*y_M_*) and define ||*y*_0_, *x_i_*|| = 0 for all *x_i_*, the minimum distance Dist(*X*[*t_s_* : *t_e_*], *Y*) could be derived as follows:

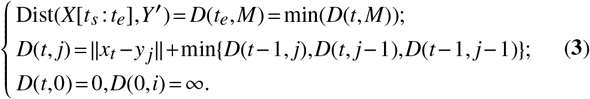

After calculating the entire distance matrix, the optimal mapping path could be traced from the end point *t_e_* in *D*(*t, M*) to the start point *t_s_*. Thus, the time complexity of this algorithm is reduced to *O*(*MN*). However, it should be noted that sDTW achieves the efficiency acceleration by giving up counting the gaps in the alignment, which makes it infeasible to the local genome-to-signal search problem in nanopore sequencing due to an order of magnitude difference in the sampling speed. A possible solution is to resample the raw signals first and then solve the alignment problem with a multi-scale scheme.

### An example of local genome-to-signal search in nanopore raw signals

Here we design a simple experiment to show the effects of different strategies on subsequence inquiry of nanopore signals.

As the raw signals have an average 8 to 9 times of redundant sampling rate (Rang *et al.*, 2018), we use the FIR (finite impulse response filter) resampling technique (Saramaki and Bregovic, 2002) to generate a 8-times compressed signal sequence *X*′ from *X* (a brief introduction of FIR resampling is given in Section S1). A query sequence with 1000 nucleotides (*Y*) and a nanopore raw signal sequence with ~ 100000 time points (*X*), which contains the query sequence, are selected to demonstrate the results (Fig. 2).

**Figure 2.**
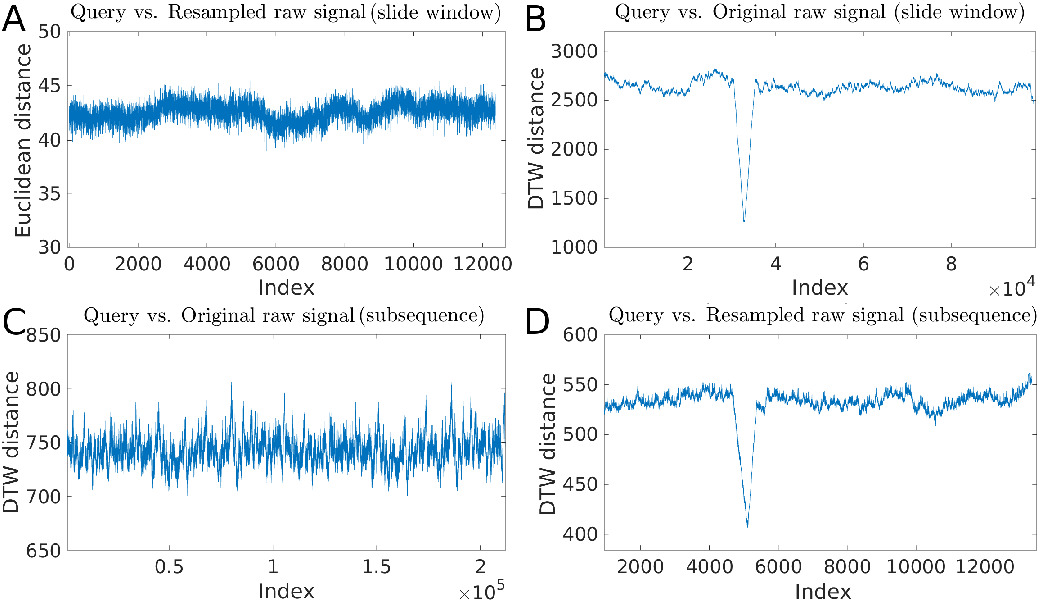
An example showing the distinguishing power of different distance measures and different variants of DTW on subsequence search. (A) Subsequence search by the Euclidean distance between the query *Y* and the resampled signal sequence *X*′; (B) Subsequence search by the DTW distance between the query *Y* and the raw signal sequence *X*; (C) Subsequence search by sDTW between the query *Y* and the raw signal sequence *X*; (D) Subsequence search by sDTW between the query *Y* and the resampled signal sequence *X*′.

As shown in Fig. 2(A), the Euclidean distance has no distinguishing power to identify the raw signal subsequence that is similar to the query sequence. On the contrary, Fig. 2(B) shows that the DTW distance could pick up the region correctly, but the sliding window based search strategy took half an hour to get the result. The sDTW method fails to identify the region (Fig. 2(C)), which is due to the massive amount of redundant sampling in the raw signals. Finally, Fig. 2(D) shows that sDTW is able to detect a sharp peak in the resampled signal sequence.

## MATERIALS AND METHODS

Two novel algorithms are proposed for direct subsequence searching and mapping in nanopore raw signals, including the direct subsequence DTW algorithm, DSDTWnano, and its accelerated algorithm, cwSDTWnano.

### Direct subsequence dynamic time warping for nanopore raw signal search

The main difficulty to apply subsequence DTW on the nanopore raw signal data is the scale difference between the query and the raw signal sequences. We propose to resolve this issue by resampling the raw signal sequence first, aligning the resampled signals to the query, remapping the warping path of the resampled signals to the original ones, and finally refining it by constrained DTW. Because the highly similar regions will result in a sharp peak, an early stop condition could be introduced to save runtime when we calculate the DTW distance along the nanopore raw signal.

We thus propose a novel algorithm, DSDTWnano (Algorithm 1), where DSDTW(·) is the subsequence dynamic time warping with an early stop condition, Resampling(·) is the FIR resampling to compress the nanopore raw signals (Saramaki and Bregovic, 2002), PathTrackback(·) is a function that recursively searches the match paths between *X*’ and *Y* that starts from *t_e_*, ReMapIndex(·) is the context-dependent constraint generation from a coarse path *W_coarse_* with a window size of *r*, cDTW(·) is the constrained dynamic time warping (Ratanamahatana and Keogh, 2005) and *s_base_* is the estimation of raw signal’s sampling rate. Because the subsequence DTW has the complexity of *O*(*MN*′) and the constrained DTW has the complexity of *O*(*rM*) (*N*′ ≈ *N*/*s_base_* is the length of resampled signals, and *N* is the length of signal *X*), the overall complexity of DSDTWnano is 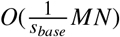.

#### Algorithm 1 DSDTWnano

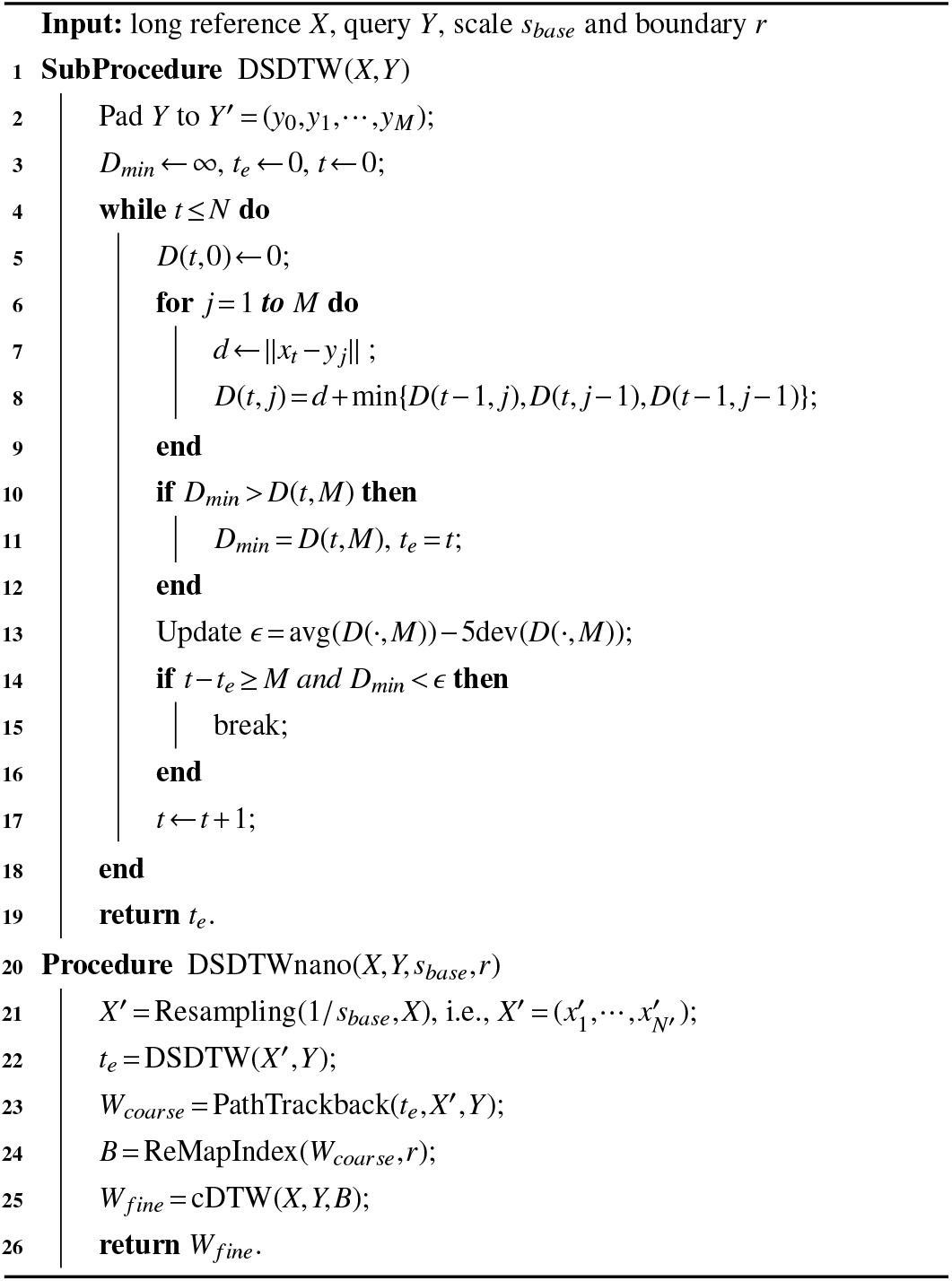

### Continuous wavelet subsequence DTW for nanopore raw signal search

Though DSDTWnano has a dramatic improvement compared with the naive sliding-window based DTW, it is still not efficient enough when handling very long raw signal sequences. To accelerate the efficiency while keeping the effectiveness, we propose cwSDTWnano, which includes several techniques to further speed up the subsequence similarity search:seeding, pre-filtering, and multi-scale search.

cwSDTWnano starts from seed search on the resampled raw signals. Based on the mapping paths of the seeds, the signal sequences with no high-similarity segment (i.e., non-hit signals) are filtered out. For the candidate signal sequences that pass the filter, a low-resolution wavelet transform is imposed on the long nanopore signal and the query signal sequences to highly compress the information, which is utilized to generate the coarse path with the help of seeds. Finally, with the multi-scale analysis of CWT, the mapping path between the query signal sequence and the raw signal sequence is calculated recursively from a lower-resolution projection to a higher-resolution one.

#### Seeds with minimal length

In genomic read mapping, the *k*-long subsequences (i.e., *k*-mers) in a query sequence are often used as a quick indicator of whether and where the reference contains the query. These k-mers are called ‘seeds’ and their inquiry is usually done through hashing. Because of the high noise and non-stable sampling rates in nanopore sequencing, it is difficult to build such a *k*-mer hash function. However, we still can use the idea of ‘seeding’ to quickly determine the range where the query signal locates in the raw signals.

One of our observations is that a query signal could be detected without ambiguity if it exceeds a certain length. Here, this certain length is denoted as the *minimal length*. An experiment is presented to show how the length of the query affects the similarity search. As shown in Fig. 2, the subsequence in the resampled raw signals with the highest similarity to the query signal will result in the minimum DTW distance, which behaves as a sharp peak. Fig. 3 shows that a 32bp- or 64bp-long query signal cannot determine a unique result because there is no distinguishable peak of the DTW distance. On the contrary, a query with 96bp or 128bp length is able to detect a clear sharp peak. Two reasons may explain why a very short query fails: (i) the noise in the raw signal degenerates the DTW distance of the true hit, and (ii) there exist multiple similar subsequences in the raw signal sequence.

**Figure 3.**
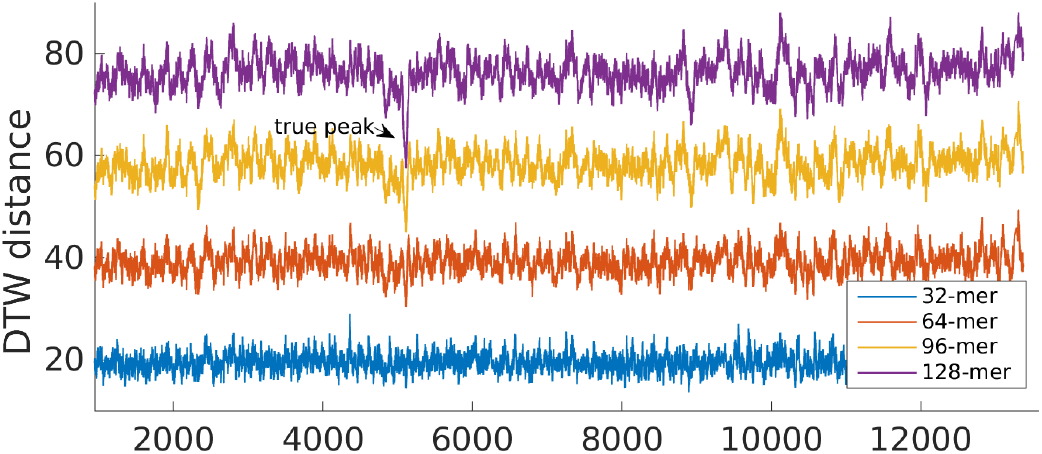
The change of the DTW distance with different query lengths.

We make comprehensive experiments, and the results prove that a length of 128bp is enough for a short query to be detected in the raw signals. Especially, if the distribution of the electrical current values in raw signals is given, it is possible to infer the theoretical *minimal length* from the given distribution, which shows the existence of the *minimal length* in any nanopore system (a brief proof is given in Section S2). Therefore, we denote a short segment in a query signal of length at least *minimal length* as a *seed*.

#### Filtering non-hit signals by seeds

Given a long query signal sequence (≥ 1000), it is possible to utilize the *seeds* to filter raw signals with no high-similarity segment, which will significantly reduce the total query time. The key observation is that if a query sequence has a highly similar region in the resampled raw signal sequence, linearly ordered seeds on the query sequence will also have a linear relationship to the hit regions in the resampled signal sequence (Fig. 4). On the contrary, if the reference sequence does not have a highly similar region to the query sequence, no linear-ordered seeds will be detected. Based on this observation, a filtering operation is developed to quickly exclude those non-hit signals:

1. Select a set of segments {*Q_i_*}_*i*=1,⋯,*K*_ from the query signal *Y* as the *seeds*;
2. For each *seed*, search in the resampled signal sequence *X*′ by sDTW(·) to get the local mapping;
3. Trackback from the endpoint of the mapping to get the mapping path of each *seed*;
4. Make a linear regression based on the mapping paths of these *seeds* and check their consistency;
5. If the consistency is weak, stop the process.

**Figure 4.**
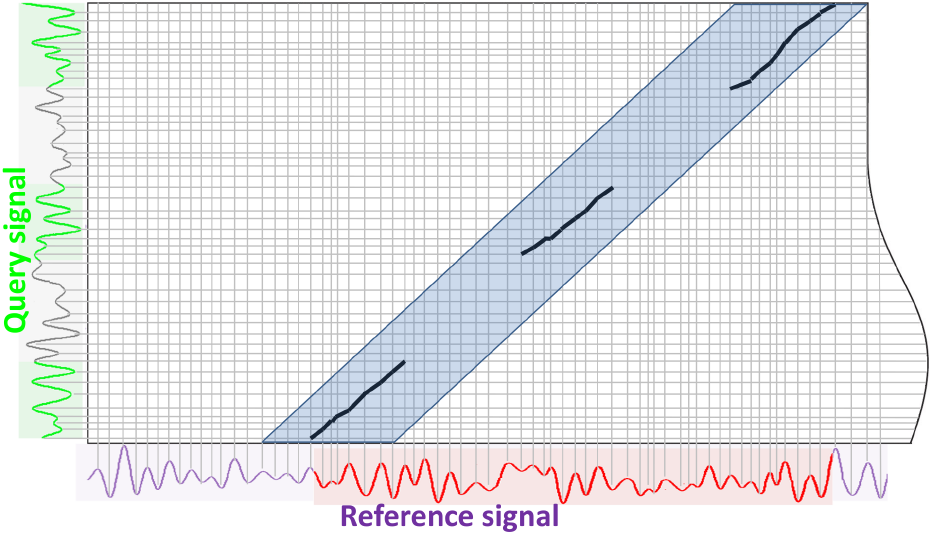
Linear relationship of the mapped path of 3 short seeds that are extracted from a long query sequence. In the figure, the seeds are marked by green color (y-axis) and the query result is labeled by red color (x-axis). It can be found that the mapped path of the seeds follows a linear relationship.

If the linear relationship of the seeds is violated, we can stop the search process to save time. For *K* seeds with length *L*, the total cost for a raw signal sequence with *N* time points is 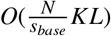, in which both *K* and *L* are very small.

#### Multi-scale search based on CWT

When handling long signal sequences, multi-scale analysis has been widely used to reduce the runtime (Salvador and Chan, 2007; Prätzlich *et al.*, 2016), and continuous wavelet transform (CWT) has been adopted to preserve the feature information (Skutkova *et al.*, 2015; Han *et al.*, 2018). Here we further combine CWT with the multi-scale analysis (Han *et al.*, 2018) and apply it to the genome-to-signal subsequence search problem.

##### Continuous wavelet transform

A continuous wavelet transform (CWT) is a formal tool that provides an overcomplete representation of a signal. In particular, the CWT of a onedimensional signal *X*(*t*) at a scale 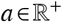 and translational value 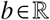, denoted as *X_a,b_*, is expressed by the following integral:

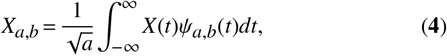

where 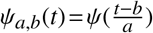 is the mother wavelet which is a continuous function in both the time domain and the frequency domain. In our algorithm, the Mexican hat wavelet is the default option, but other wavelet functions are also applicable (Torrence and Compo, 1998).

##### Multi-scale representation

For the convenience of analysis, we fix the translational value *b* as the same index correspondence as *X*. That is, the transformed signals (spectrum) have the same length and retain peer-to-peer index to *X*. Here we use CWT(*X, a*) to denote the transformed spectrum of *X* with the scale parameter *a*. A feature extraction procedure can be carried out (denoted as PickPeaks(·)) to reduce the length of a signal *X*: (i) obtain the spectra CWT(*X,a*); (ii) normalize CWT(*X,a*) based on Z-score normalization; (iii) extract peaks and nadirs from each spectrum as the feature sequence. The length of a signal could be dramatically reduced by more than *a* times for a classic nanopore raw signal sequence.

##### Coarse path generation

As introduced before, a number of *seeds* are used and their mapping paths with the resampled signal sequences are recorded. These short mapping paths can be used as anchors in the construction of the coarse mapping path between the query sequence and the raw signal sequence using the lowest resolution transform (i.e., with maximal level coarsening scale) from CWT:

1. Given the query sequence *Y* with length *M*, get the maximal level coarsening scale *a* = log_2_(*M*) − 2;
2. Get the feature signals for both CWT(*X*′,*a*) and CWT(*Y,a*);
3. Run the subsequence DTW on the feature signals and get all the paths;
4. Find out the coarse path that covers the seeds;
5. Combine both the seeds and the coarse path to generate a more detailed path.

Then, the generated coarse mapping path is fed into cwDTW (Han *et al.*, 2018) to determine the final mapping.

##### The continuous wavelet subsequence DTW

Algorithm 2 shows cwSDTWnano, where cwDTW(·) is the continuous wavelet-based multi-level DTW (Han *et al.*, 2018), SelectSeeds(·) is the procedure to get *K* segments with length *L* from *Y*, CheckFalse(·) is the filtering of false alignment described in Section 26, ReMapIndex(·) is the context-dependent constraint generation from a coarse path *W_coarse_* with a window size of *r*, CWT(·) is the continuous wavelet transform and PickPeaks(·) is the procedure to get feature sequence (Han *et al.*, 2018), CoarsePath(·) is the coarse path generation procedure described in the previous paragraph and cDTW(·) is the constrained DTW (Ratanamahatana and Keogh, 2005). We notice that the false filtering procedure has a complexity of 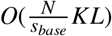 and the procedure of cwDTW(·) is bounded within *O*(*N*log*N*). Thus the overall complexity for Algorithm 2 is 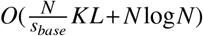, which has an obvious advantage when the signal length increases.

#### Algorithm 2 cwSDTWnano

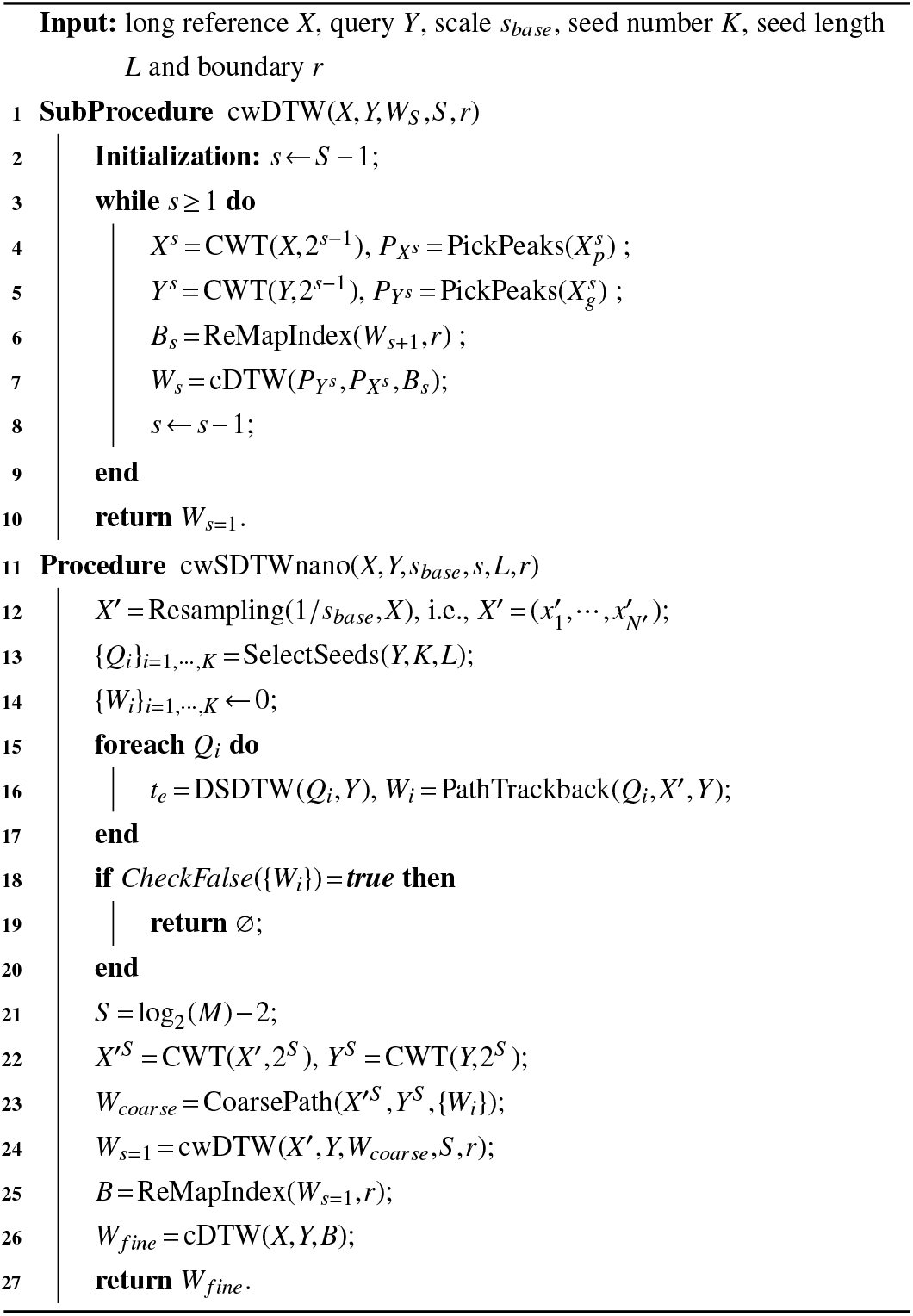

## EXPERIMENTS AND RESULTS

### Datasets

Three real-world nanopore sequencing datasets are used in our experiments, among which the first (human) and second (lambda phage) are used to evaluate the accuracy of our proposed local search algorithms. The third dataset (E. coli) is used to show the power of our algorithms on the discrimination of hit and non-hit signals, as well as the low-coverage SNP detection.

The first dataset is a subset of the publicly available human data, which comes from human chromosome 21 from the Nanopore WGS Consortium (Jain *et al.*, 2018) and contains 6318 sequenced reads. The samples in this dataset were sequenced from the NA12878 human genome reference on the Oxford Nanopore MinION using 1D ligation kits (450 bp/s) with R9.4 flow cells (raw signals downloaded from the nanopore-wgs-consortium http://s3.amazonaws.com/nanopore-human-wgs/rel3-fast5-chr21.part03.tar). We denote this dataset as the Human21 database.

The second and third datasets are from the genome of lambda phage and E. coli, respectively. These two datasets were all prepared and sequenced at the University of Queensland by Prof. Lachlan Coin’s lab. The lambda phage dataset contains 27004 reads and the E. coli dataset contains 27608 reads. The samples were sequenced on the MinION device with 1D protocol on R9.4 flow cells (FLO-MIN106 protocol). We denote these two datasets as the Lambda phage database and the E. coli database, respectively. Specifically, E. coli has a relatively low coverage (20×).

To comprehensively evaluate the performance of the algorithms, we created a subset by randomly sampling 3000 reads from Human21 and Lambda phage (data avaliable at https://drive.google.com/drive/folders/1LuOxg9qE1l9AuDcfyUz9aF10X4cgmX5t?usp=sharing). The average length of the DNA sequences in the sampled datasets is 7890 and 8461 for Human21 and Lambda phage, and the average length of the nanopore raw signal sequences is 65947 and 69715, respectively.

### Similarity criteria

#### Edit mapping error of a local search

Suppose the reference genome is known, we may use the edit mapping error to evaluate the difference between the mapping path generated by a local genome-to-signal search algorithm and the global mapping path.

Specifically, given a nanopore raw signal sequence, as we know the reference genome, it is possible to find the corresponding genomic region to the raw signals (Li, 2018). Therefore, the global mapping path *W*′ between the genomic region and the raw signal sequence can be derived by the original dynamic time warping (Stoiber *et al.*, 2016; Han *et al.*, 2018).

For a genomic region *G* = *g*_1_ *g*_2_⋯*g_L_* and its corresponding raw signal sequence *R* = ⊓ *r*_2_⋯*r_N_*, the accuracy of the mapping path *W* generated by a local search algorithm is defined as:

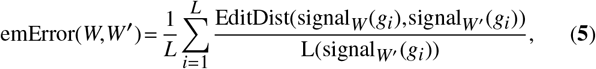

where signal*_x_*(*g_i_*) returns the set of signal indexes {*r_j_*} that corresponds to the query sequence position *g_i_* from a certain mapping path *x*∈{*W,W*′}. This is because on average, each nucleotide corresponds to 8 to 9 signals in the raw signal sequence due to the redundant sampling in nanopore. EditDist(·) is the edit distance and L(·) is the size of the signal index set.

For example, if we have a query *G* = *g*_1_*g*_2_*g*_3_ with *L* = 3. Suppose its local mapping path *W* is {(10,1), (11,1), (12,1), (13,2), (14,2), (15,2), (16,3), (17,3)}, and the global mapping path *W*′ is {(11,1), (12,1), (13,2), (14,2), (15,2), (16,3), (17,3)}. Then we will have the edit distance for *g*_1_, *g*_2_ and *g*_3_ being 1, 0 and 0, respectively, and thus emError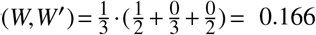. If the local search algorithm returns a perfect mapping path, the error is zero. Note that the error may exceed 100% if the mapping is way off.

#### Normalized signal distance of a local search

Suppose the reference genome is unknown or not accurate, it is difficult to obtain the global mapping. In this case, we may use the normalized signal distance (*nDist*) to evaluate the similarity between the mapped raw signal and the corresponding reference.

Given the mapping path *W* generated by a local search algorithm, the genome-to-signal similarity is defined as:

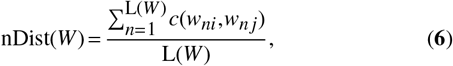

where L(*W*) is the length of the mapping path *W*, and *c*(*w_ni_*, *w_nj_*) is the absolute (or, Z-score) difference of the *n*th aligned element between the two signal points *x_i_* (the nanopore raw signal) and *y_j_* (the expected signal from the *k*-mer pore model). Different from emError(*W, W*′), here nDist(*W*) is defined over one mapping path *W* only, instead of over two mapping paths *W* and *W*′.

For a new dataset with multiple sequences, the normalized signal distance is important for the discrimination of hit and non-hit signals, as well as for data classification and clustering analysis.

### Performance

#### Visualization of a detailed example

To demonstrate the effectiveness of our algorithms in discovering the corresponding subsequences in the raw signal sequence, we give one example in Fig. 5 to show the detailed steps of local genome-to-signal search.

**Figure 5.**
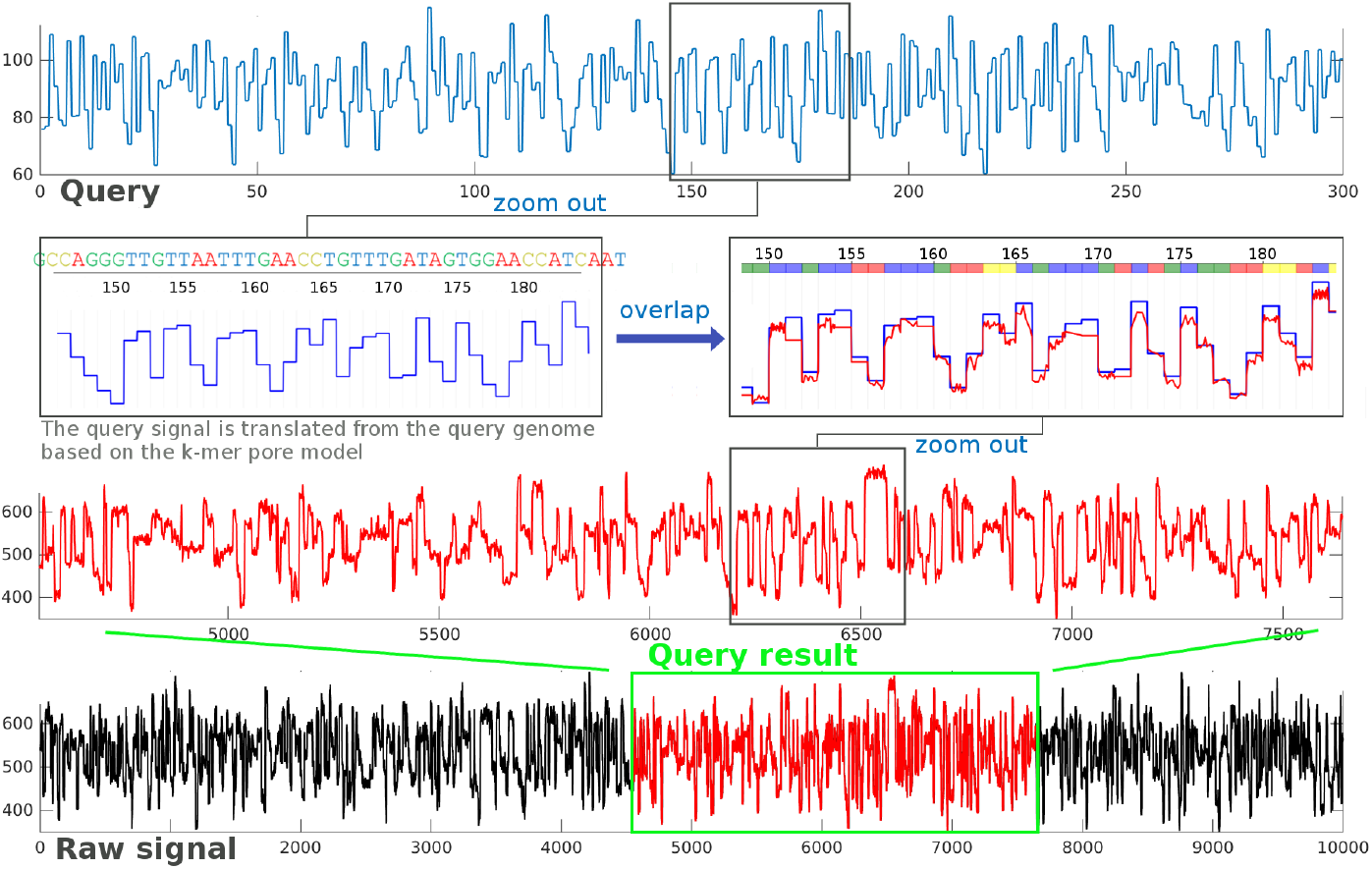
A demonstration of the raw signal similarity search and mapping by our algorithm. Here, the top signal (blue) is a query signal of length 300, and the bottom signal (black) is a nanopore raw signal of length 10000. The zoom-out subfigure locating at [145:185] shows how the query signal corresponds to the {*A,C,G,T*} nucleotides. The red signal that locates at [4553:7641] on the raw signal is the query result. By further selecting the segment [6197:6594] on the raw signal and overlapping it with the segment [145:185] on the query signal, a superimposed image with high degree of overlapping is produced.

Here, a short region of the DNA sequence with 300bp length is selected as the query sequence and a raw signal sequence with 10000 time points is served as the signal database (the black signal depicted on the bottom of Fig. 5). Both the DNA sequence and the raw signal sequence are selected from the Human21 dataset.

Below are the four steps of the genome-to-signal search procedure:

A. The query sequence is translated into a query signal sequence based on the 6-mer pore model provided by Nanopore Technologies (the blue signal sequence depicted on the top of Fig. 5).
B. Run the DSDTWnano algorithm to obtain the detailed region ([4553:7641]) on the raw signal sequence that has the highest similarity with the query signal sequence (the red signal region depicted on the bottom of Fig. 5). This query operation takes 29 ms and results in a normalized signal distance of 0.1556 between the query signal sequence and the red region of the raw signal sequence. Typically, a normalized signal distance ranging from 0~0.20 indicates a good hit.
C. By comparing the zoomed out regions of the query and raw sequences, we can find that these two signals are very similar to each other. However, it should be noted that the query result in the raw signal sequence is about 9× longer than the query signal, which is the typical difference in the sampling speed in nanopore sequencing. Nevertheless, our algorithm still produced an accurate mapping.
D. By further selecting the segment [145:185] on the query sequence and the segment [6197:6594] on the raw sequence, we may align and visualize them according to the mapping path produced by our algorithm.

#### Accuracy analysis

The performance of DSDTWnano and cwSDTWnano is evaluated using the subset of the Human21 dataset and the Lambda phage dataset. In doing so, we randomly select a segment with length *l* as the query sequence and then run the two algorithms on the corresponding raw signals to find out its maximal response mapping. Finally, we compare the query results of DSDTWnano and cwSDTWnano with the global mapping by the edit mapping error.

We first run an experiment of DSDTWnano and cwSDTWnano (with parameter *K* = 3 and *L* = 128) with the mapping boundary *r* = 50 and the query length *l* = 1000. As shown in Fig. 6(A), the distribution of the edit mapping error of cwSDTWnano is very similar to that of DSDTWnano, and the majority of the error ranges between 0 and 0.01. Fig. 6(B) shows the scatter plot between the edit mapping error of DSDTWnano (x-axis) and that of cwSDTWnano (y-axis), which indicates that most of them are the same (on the diagonal of the scatter map). The outliers of cwSDTWnano may be caused by the coarsening in the multi-scale analysis.

**Figure 6.**
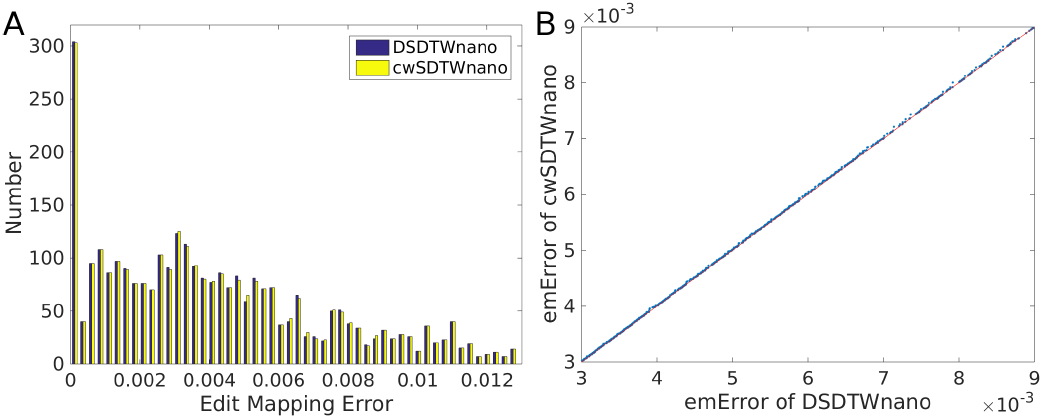
Statistics of the edit mapping error of our algorithms on the Human21 database, where for both DSDTWnano and cwSDTWnano the mapping boundary *r* = 50 and query length *l* = 1000. (A) Distribution of the edit mapping error of DSDTWnano (in yellow) and cwSDTWnano (in blue). (B) Scatter plot between the edit mapping error of the DSDTWnano (x-axis) and that of cwSDTWnano (y-axis).

We then challenge both algorithms with different lengths *l* of the query sequence and different radius *r* of the mapping path boundary. The average edit mapping error of the query results by DSDTWnano and cwSDTWnano on the Human21 database and the Lambda phage database are summarized in Tables 1 and 2, respectively. We can find that for queries with different lengths, (i) DSDTWnano almost always outputs a query result within 0.01 edit mapping error, and no larger than 0.006 for most of the cases; (ii) the edit mapping error of cwSDTWnano can also be controlled around 0.006 if a suitable *r* is selected (*r* = 50 for *l* ⩽ 2000 and *r* = 70 for *l* ⩽ 4000). This is normal as the performance of cwSDTWnano depends on the mapping boundary *r* which is required for the coarsening of the input signals. As a result, although human and lambda phage are two completely different species, from the little difference between the two tables, we know that the performance of our methods is stable and consistent over different species.

**Table 1.** The average edit mapping error on the Human21 database

| Edit Mapping Error | | $l=600$ | $l=1000$ | $l=2000$ | $l=3000$ | $l=4000$ |
| --- | --- | --- | --- | --- | --- | --- |
| DSDTW<br>nano | $r=30$ | 0.003992 | 0.004477 | 0.005322 | 0.005602 | 0.005575 |
| | $r=50$ | 0.004131 | 0.004440 | 0.005213 | 0.005361 | 0.005326 |
| | $r=70$ | 0.004092 | 0.004533 | 0.005147 | 0.005368 | 0.005158 |
| cwSDTW<br>nano | $r=30$ | 0.004444 | 0.007988 | 0.012675 | 0.019269 | 0.030104 |
| | $r=50$ | 0.004183 | 0.004651 | 0.005647 | 0.005867 | 0.006013 |
| | $r=70$ | 0.004100 | 0.004598 | 0.005308 | 0.005504 | 0.005395 |

**Table 2.** The average edit mapping error on the Lambda phage database

| Edit Mapping Error | | $l=600$ | $l=1000$ | $l=2000$ | $l=3000$ | $l=4000$ |
| --- | --- | --- | --- | --- | --- | --- |
| DSDTW<br>nano | $r=30$ | 0.003813 | 0.003940 | 0.004631 | 0.004898 | 0.004686 |
| | $r=50$ | 0.003544 | 0.004059 | 0.004527 | 0.004667 | 0.004583 |
| | $r=70$ | 0.003763 | 0.003933 | 0.004347 | 0.004453 | 0.004265 |
| cwSDTW<br>nano | $r=30$ | 0.004674 | 0.006521 | 0.014902 | 0.035661 | 0.052850 |
| | $r=50$ | 0.003791 | 0.004549 | 0.005384 | 0.005728 | 0.005973 |
| | $r=70$ | 0.003791 | 0.004294 | 0.004673 | 0.004758 | 0.004667 |

Because cwSDTWnano has two extra parameters, *K* and *L*, to define the seed number and the seed length, we further analyze the parameter sensitivity. Table 3 summarizes the average edit mapping error on the Human21 database for cwSDTWnano with different seed numbers *K* and seed lengths *L* (here the search radius *r* is set to 50). From Table 3 we can find that the seed length has an influence on the quality of the result but the number of seed does not have. Also, the edit mapping error demonstrated in Table 3 indicates that cwSDTWnano is robust for *K* ⩾ 3 and *L* ⩾ 128. Thus a very short seed may cause false dismissals, whereas a seed length of 128 can ensure the correctness of cwSDTWnano for almost all the queries, and a seed length of 192 is sufficient for a dataset with raw signals with reasonably good quality.

**Table 3.** The average edit mapping error of query results on the Human21 database for cwSDTWnano with different configurations

| Edit Mapping Error | $l=600$ | $l=1000$ | $l=2000$ | $l=3000$ | $l=4000$ |
| --- | --- | --- | --- | --- | --- |
| $K=3, L=128$ | 0.004183 | 0.004651 | 0.005647 | 0.005867 | 0.006013 |
| $K=4, L=128$ | 0.004155 | 0.004819 | 0.005569 | 0.006062 | 0.005725 |
| $K=5, L=128$ | 0.004202 | 0.004643 | 0.005549 | 0.005785 | 0.005832 |
| $K=3, L=192$ | 0.004444 | 0.004626 | 0.005355 | 0.006047 | 0.005624 |
| $K=4, L=192$ | 0.004223 | 0.004745 | 0.005398 | 0.005949 | 0.005759 |
| $K=5, L=192$ | 0.004177 | 0.004950 | 0.005322 | 0.006114 | 0.005730 |

#### Runtime analysis

For a database with a number of raw signals, the running time for a query is also important. Generally, the runtime of DSDTWnano is about 450 ms and that of cwSDTWnano is about 200 ms for a query sequence with 1000bp in length on a 100000 time points raw signal sequence. When the query length grows, the runtime may increase considerably if there are hundreds or thousands of raw signals. Under this condition, cwSDTWnano is suitable because it can accelerate the query process remarkably by a multi-scale strategy. In this subsection, the runtime for both DSDTWnano and cwSDTWnano is investigated.

Fig. 7 demonstrates the runtime of our algorithms. All the execution time is collected on a Fedora25 system with 128Gb memory and two E5-2667v4 (3.2 GHz) processors. From Fig. 7(A) we can find that DSDTWnano has a much higher execution time compared with cwSDTWnano when the query length increases, whereas cwSDTWnano keeps a low computational cost. Specifically, the runtime of cwSDTWnano is always shorter than 900 ms even searching a 6000bp-long query on a raw signal with 2 × 10^5^ time points. From Fig. 7(B) we can find that the runtime of cwSDTWnano does not exceed 1500 ms when searching a 1000bp-long query on a raw signal with 1 × 10^6^ time points.

**Figure 7.**
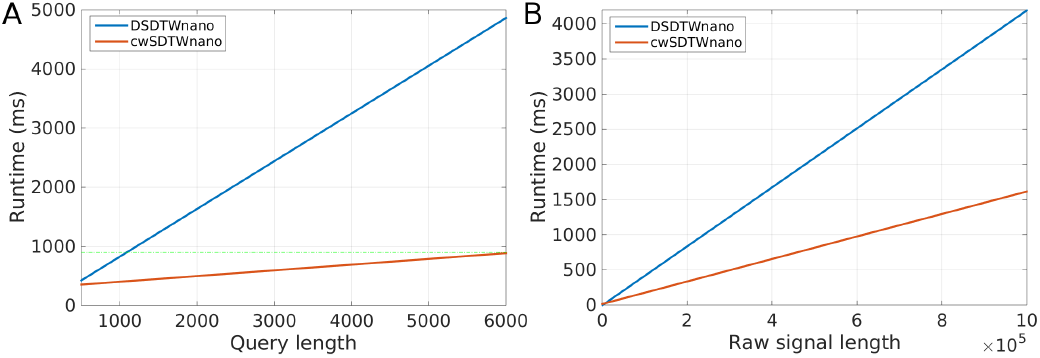
The runtime of our algorithms with different query lengths and raw signal lengths (r = 50, *K* = 3 and *L* = 128). (A) The runtime of DSDTWnano and cwSDTWnano on a 2 × 10^5^-long raw signal sequence when the length of the query changes; (B) The runtime of DSDTWnano and cwSDTWnano for a 1000bp-long query when the length of raw signals changes.

In practice, we recommend to run DSDTWnano if the query length is short, and run cwSDTWnano otherwise.

#### Discrimination of hit and non-hit signals

A fundamental task in nanopore sequencing is that, given a query sequence and a raw signal database, whether we can find a set of signal segments (subsequences of raw signals) that are similar to the query, i.e., distinguishing the hit signals from the non-hit ones. This is necessary because in some applications such as SNP detection, the task is to find some non-standard signals, in which multiple numbers of hit signals are required for the reliable detection of these events. However, it is not easy to implement a scoring function to discriminate hit and non-hit signals, especially when the reference genome is unknown or not accurate. Here, we show that the normalized signal distance (*nDist*) is a good similarity measurement in such a task.

We use the E. coli database to test the performance of *nDist* with DSDTWnano to identify hit and non-hit signals. To construct the benchmark, we randomly select 600bp-long and 1000bp-long subsequences from the E. coli genome as the query sequences, each with 200 samples. For each raw signal sequence in the E. coli database, as its corresponding reference sequence is known, we are able to get the true label of each sequence. Since the sequencing coverage of the E. coli dataset is around 20, we use all the hit signals as the *true* set, and randomly sample 200 non-hit as the *false* set. For each pair of the query sequence and the raw signals in either the *true* or *fasle* set, we run DSDTWnano to obtain the local mapping path and the corresponding *nDist* score.

As shown in Fig. 8, we observe that (i) there are two well-separated distributions of *nDist*, where almost all the left (right) belongs to the *true (fasle)* set; and (ii) *nDist* from the query sequence with different lengths reside in the same distribution. Thus, *nDist* can distinguish the *true* set (i.e., hit signals) from the *false* one (i.e., non-hit signals) regardless of the query length. It is obvious that there is a clear boundary around *nDist =* 0.2 that could separate the *true* and *false* sets, which is used as the threshold in practice. Specifically, such a characteristic of *nDist* naturally constructs a linear classifier for further data classification task.

**Figure 8.**
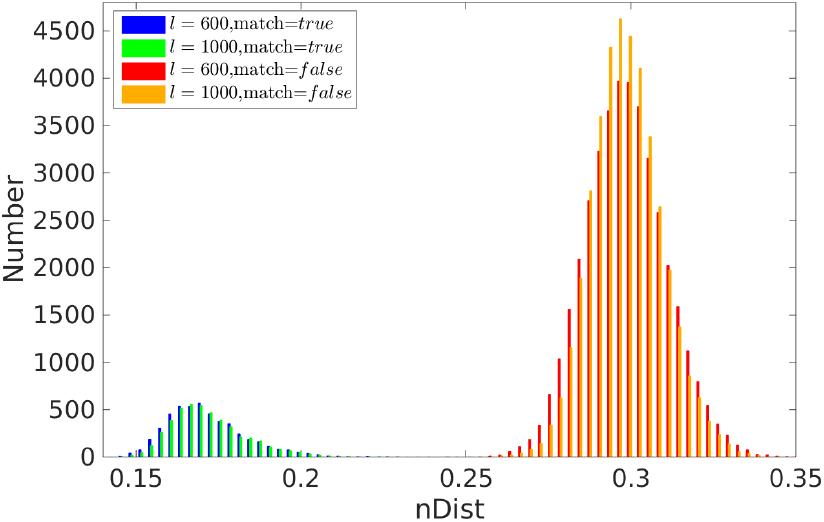
The histogram of the normalized signal distance (*nDist*) on the E. coli database, which is generated from 200 query sequences whose length is 600bp-long or 1000bp-long. Each query sequence has 200 *non – hit* raw signals (denoted as the *false* set) and about 20 *hit* raw signals (denoted as the *true* set).

#### Comparison with read-based approach

As discussed in introduction, there exist two subsequence inquiry approaches for nanopore-based targeted sequence analysis: signal-based and read-based. In the signal-based approach, the genomic region of interest will be first translated into the expected signals by the k-mer pore model, and then be inquired as signal against the database containing the raw signals. In the read-based approach, the raw signals are first transformed into the reads by base-calling, and then the genomic sequence of interest will be used for detecting the similar subsequences within these reads. Our proposed algorithms belong to the former, while a variety of standard read mappers (e.g., minimap2 (Li, 2018), BLAST (Altschul *et al.*, 1997), and others (Sedlazeck *et al.*, 2018; Sovic *et al.*, 2016; Langmead and Salzberg, 2012; Li and Durbin, 2010)) belong to the latter. A natural question to ask is if there really exists an advantage of the signal-based approaches over the read-based methods in processing nanopore sequencing data.

To answer this question, we use our in-house tool DeepSimulator (Li *et al.*, 2018) to simulate 20,000 reads and signals at two typical sequencing accuracy (say, 80% and 90%) from a given 1M bp genomic region that encompasses Human DGCR8 gene (essential for microRNA biogenesis (Wang *et al.*, 2007)). Then we randomly select subsequences at different lengths (say, 200 bp, 400 bp, and 800 bp) within this 1M bp region, each with 5 samples, as the query sequences to perform subsequence inquiry. The programs to compare with our DSDTWnano are minimap2 and BLAST, which are processed with default parameters.

As the ground-truth is known during simulation, for each subsequence, we denote those reads that fully contain (not contain) this subsequence as hit (non-hit) reads. In order to eliminate ambiguity, we exclude those reads that overlap with this subsequence. Thus, for each method, the purpose is to identify as much as hit reads as possible, while avoiding classifying those non-hit reads as hits.

For our DSDTWnano method, it is straightforward to distinguish hit and non-hit reads by setting the threshold as *nDist* = 0.2. However, for minimap2 and BLAST, it is not straightforward to do so as they will report some reads with low-similarity or low-quality. To remove them, we set a length of alignment (LALI) threshold 0.75 · *L* for BLAST and minimap2 where *L* is the length of the query subsequence. For example, if the length of a subsequence is 400 bp, then we exclude those reads whose LALI is below 300.

We measure the success rate of subsequence inquiry in the following terms: balanced accuracy (bAcc), precision (Prec), sensitivity (Sens), specificity (Spec), and Matthews correlation coefficient (Mcc). In order to calculate these terms, we define the True Positives (TP) and True Negatives (TN) as the numbers of correctly identified hit and non-hit reads, respectively, where False Positives (FP) and False Negatives (FN) are the numbers of misclassified hit and non-hit reads, respectively. Precision, sensitivity and specificity are defined as *TP*/(*TP* + *FP*), *TP*/(*TP* + *FN*) and *TN*/(*TN* + *FP*), respectively. Balanced accuracy is the average of sensitivity and specificity.

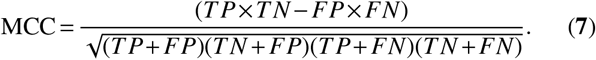

As shown in Table 4 and Table 5, our method outperforms BLAST and minimap2 by a large margin, especially when the length *L* of the query subsequence is short, regardless of the dataset with a relatively low or high sequencing accuracy. Specifically, for *L* = 200 at 80% sequencing accuracy, our method achieves 0.947 Mcc, 0.913 sensitivity, 0.914 precision and 0.948 balanced accuracy, respectively, which are 58.5%, 66.5%, 4.6% and 33% higher than minimap2, and 26.2%, 43%, 27.2% and 7.7% higher than BLAST; for 90% sequencing accuracy, our method achieves 0.978 Mcc, 0.985 sensitivity, 0.909 precision and 0.992 balanced accuracy, respectively, which are 25.5%, 19.8%, 8.2% and 13.9% higher than minimap2, and 14.5%, 2.6%, 17% and 8% higher than BLAST.

**Table 4.** Performance of subsequence inquiry on the simulated dataset at 80% sequencing accuracy

|  |  | bAcc | Prec | Sens | Spec | Mcc |
| --- | --- | --- | --- | --- | --- | --- |
| $L=200$ | Minimap2 | 0.618 | 0.873 | 0.248 | **0.995** | 0.362 |
|  | BLAST | 0.871 | 0.642 | 0.483 | 0.932 | 0.685 |
|  | Our proposal | **0.948** | **0.914** | **0.913** | 0.986 | **0.947** |
| $L=400$ | Minimap2 | 0.783 | 0.992 | 0.534 | 0.984 | 0.623 |
|  | BLAST | 0.923 | 0.953 | 0.872 | 0.978 | 0.927 |
|  | Our proposal | **0.956** | **0.996** | **0.942** | **0.993** | **0.964** |
| $L=800$ | Minimap2 | 0.947 | 0.922 | 0.791 | **0.997** | 0.821 |
|  | BLAST | 0.934 | 0.955 | 0.913 | 0.975 | 0.944 |
|  | Our proposal | **0.991** | **0.996** | **0.981** | **0.997** | **0.989** |

**Table 5.** Performance of subsequence inquiry on the simulated dataset at 90% sequencing accuracy

|  |  | bAcc | Prec | Sens | Spec | Mcc |
| --- | --- | --- | --- | --- | --- | --- |
| $L=200$ | Minimap2 | 0.853 | 0.827 | 0.787 | 0.941 | 0.723 |
|  | BLAST | 0.912 | 0.739 | 0.959 | 0.933 | 0.833 |
|  | Our proposal | **0.992** | **0.909** | **0.985** | **0.998** | **0.978** |
| $L=400$ | Minimap2 | 0.942 | 0.991 | 0.924 | **0.999** | 0.925 |
|  | BLAST | 0.949 | 0.932 | 0.918 | 0.983 | 0.926 |
|  | Our proposal | **0.999** | **0.999** | **0.999** | **0.999** | **0.999** |
| $L=800$ | Minimap2 | 0.935 | 0.903 | 0.921 | 0.985 | 0.914 |
|  | BLAST | 0.937 | 0.946 | 0.967 | 0.982 | 0.951 |
|  | Our proposal | **0.999** | **0.999** | **0.999** | **0.999** | **0.999** |

### Case study

Two case studies of SNP detection and haplotyping classification are presented to demonstrate the application of our algorithms in targeted sequencing (see Fig.9).

#### SNP detection

Detecting genetic variations, such as single nucleotide polymorphisms (SNPs), in a specific region of the genome is a major task in targeted sequencing. Currently, the identification of SNPs is mainly done by resequencing approach (i.e., searching for differences between aligned reads and the reference genome) or assembly approach (i.e., *de novo* assembling consensus read sequences against a reference genome) (Magi *et al.*, 2017). Recently, a few studies explored the capability of nanopore sequencing to identify SNPs (Quick *et al.*, 2016), which conclude that to reach a high detection rate (such as ~ 90%), more than 60× sequencing coverage is needed (Jain *et al.*, 2015).

**Figure 9.**
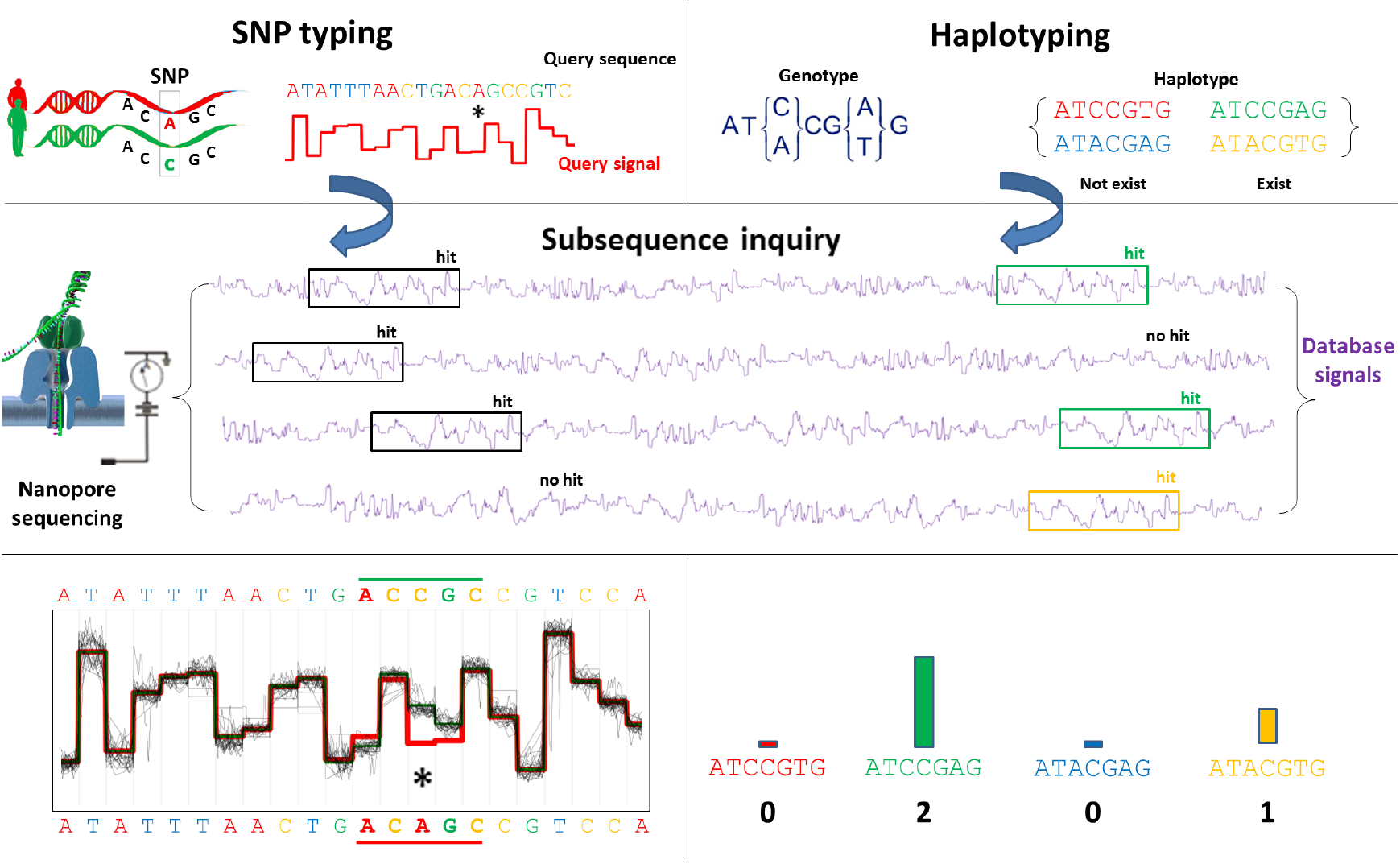
Two use cases to demonstrate the potential applications of the reference-to-signal pipeline in nanopore-based targeted sequencing. Left: SNP typing. Here the query sequence is the SNP-containing genomic region. The aligned raw signals upon the query sequence with a SNP (here is *A* → *C*) is shown in the bottom. A clear difference between the raw signals (black squiggle curves) and the expected signals (red and green curves for reference and mutated sequences, respectively) indicates strong evidence of SNP. Right: haplotyping. Here the query sequences are the four candidate haplotypic sequences. A raw signal should be classified to the most similar haplotype if it passes the similarity threshold. The count of each haplotype is shown in the bottom.

A case study is presented to demonstrate how we can identify and visualize SNPs based on the nanopore raw signals at a low sequencing coverage on a targeted genome region. The experiment is carried out on the E. coli dataset with a series of relatively low coverage (10×, 15× and 20×) and a number of randomly mutated SNPs (10,100,1000 and 10000 SNPs) on the genomic region covering the first 2.5Mbp. Here we choose 2.5Mbp because this length is roughly the upper bound of the targeted sequencing reported so far using the CATCH (Cas9-assisted targeting of chromosome segments) technology (Bennett-Baker and Mueller, 2017).

In doing so, we first generate a mutated genome by randomly substituting *n* bases on the reference genome. Then we randomly select *N* raw signals from the signal database to fit the required coverage *c*. Afterwards, given a mutated genome with *n* SNPs and the signal database at coverage c, we extract 600bp-long sliding window sequences with a step size of 300bp from the mutated genome and use them as queries in cwSDTWnano, to locate the candidate raw signal segments and positions that might contain a SNP. The SNP positions are then detected based on the mismatches between the aligned signals and the expected signals of the reference sequence (without mutation), as measured by Z-score. After the candidate SNP regions are detected, for each position within this region, four mutated sequences each with that position being {*A,C,G, T*}, respectively, are used as the query to search against the signal database. Finally, the mutation with the expected signal closest to the observed signals in the database is chosen as the detected SNP at the candidate position (more details are given in Section S3).

To evaluate the performance of our algorithms in the low coverage situation, we calculate the SNP detection rate and compare our method with Nanopolish (Quick *et al.*, 2016) at different coverages *c* and different SNP numbers *n*. Table 6 summarizes the experimental results with different numbers of SNPs and different signal coverages. Our method always dramatically outperforms Nanopolish, especially at low coverage. This is an important feature for nanopore-based targeted sequencing because nanopore does not need PCR amplification and thus often has a low coverage, especially for single cell experiments. When the coverage is as low as 10, Nanopolish almost fails to detect any SNPs, whereas our method can detect roughly half of them.

**Table 6.** The SNP detection ratio under different signal coverages

| SNP detection rate | | $n = 10$ | $n = 100$ | $n = 1000$ | $n = 10000$ |
| --- | --- | --- | --- | --- | --- |
| Our method | $c = 10$ | 0.500 | 0.480 | 0.530 | 0.510 |
| | $c = 15$ | 0.800 | 0.820 | 0.795 | 0.803 |
| | $c = 20$ | 0.900 | 0.890 | 0.889 | 0.897 |
| Nanopolish | $c = 10$ | 0.000 | 0.040 | 0.051 | 0.044 |
| | $c = 15$ | 0.400 | 0.440 | 0.431 | 0.447 |
| | $c = 20$ | 0.700 | 0.780 | 0.771 | 0.769 |

An example of SNP identification is shown in Fig. 10, which is a region of aligned raw signals (a full mapping can be found in Section S3). Here the red (green) curves indicate the 6-mer pore model for the reference (mutated) sequence centered at the candidate SNP position. The aligned nanopore signals are shown in black. There is a clear difference of the pore model at the SNP position, which indicates a strong evidence.

**Figure 10.**
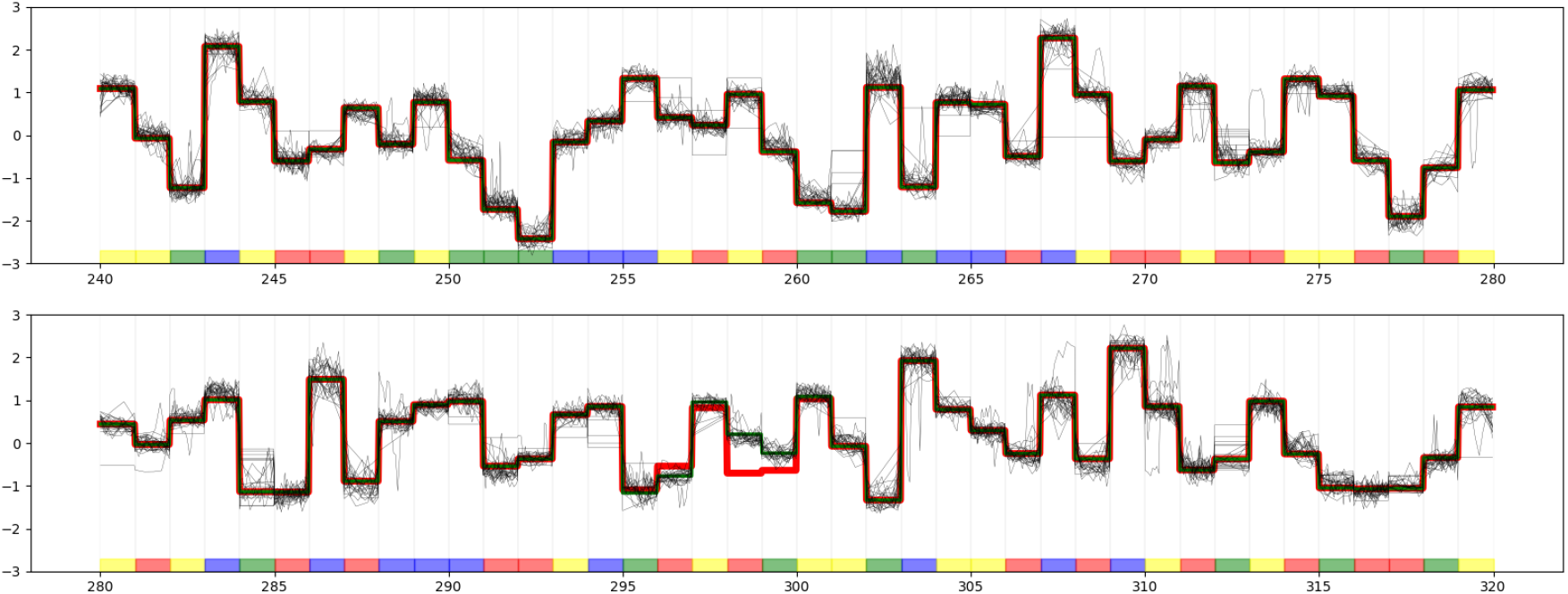
Illustration of SNP detection by cwSDTWnano. A, C, G and T on the reference sequence (query) are labeled in red, yellow, green and blue, respectively. The aligned nanopore signals are shown in the black squiggle curves and the red (green) curves indicate the 6-mer pore model for the reference (mutated) sequence centered at the candidate SNP position.

In summary, experiments on the E. coli dataset demonstrate that accurate SNP detection (around 90%) can be achieved by a low coverage (i.e., 20×) with the help of our algorithms. The success of our algorithms lies in two folds: (i) the signal-level operation reserves more information, and (ii) the normalized signal distance measurement effectively filters out non-hit signals and identifies hit signals.

#### Haplotype classification

The genome of a lot of eukaryotic species, including human, is diploid. Each of its autosomes (i.e., non-sex chromosome) comes in two copies. These parental copies are affected by different SNPs, and the assignment of these SNPs to each copy is defined as haplotyping (Consortium *et al.*, 2005). Currently, there are two major approaches to perform haplotype classification: (i) statistical methods, which assume that the haplotypes to be computed are a mosaic of reference haplotype blocks that arise from recombination during meiosis, and use maximum-likelihood estimation to solve the problem (Browning and Browning, 2011); and (ii) sequencing approach, which addresses the haplotype classification directly from the sequencing reads (Patterson *et al.*, 2015).

With the rise of targeted sequencing techniques, the haplotyping within a selected genomic region becomes possible. Here, we formulate the targeted haplotyping problem as searching all the possible haplotypic sequences within a selected genomic region against the raw nanopore signals. As nanopore data with known haplotyping are not available, we use our inhouse tool DeepSimulator (Li *et al.*, 2018) to simulate signals and reads at a relatively low sequencing accuracy.

In particular, we generate two haplotypes of the 42 kb human MDM2 oncogene centered in a 200 kb genomic region. The MDM2 protein is a ubiquitin ligase that plays a critical role in regulating the levels and activity of the p53 protein (Atwal *et al.*, 2007). The two SNPs that we choose to generate the two haplotypes locate at positions 285 C/G and 309 T/G, which are shown to be associated with an earlier age of tumor onset (Renaux-Petel *et al.*, 2014).

The experiment is conducted as follows: (i) the two assigned haplotypes in our simulation are 285C-309T and 285G-309G, respectively; (ii) the coverage of simulated signals/reads (the average accuracy of the simulated reads is about 85%) in this 200 kb genomic region is about 20× for each haplotype; (iii) four sequences with 800 bp length that cover this haplotype region (say, 285C-309T, 285C-309G, 285G-309T, and 285G-309G) are used as query, to find out the segments of the raw signals that cover this 800 bp region (denoted as hit); (iv) for each hit signal, the normalized signal distance (nDist) of these four sequences are calculated and the minimum one is selected as the haplotyping label.

Among the ~ 5000 generated signals/reads, 44 of them are hit signals that cover this 800 bp haplotype region. As the ground-truth of the haplotype for each hit signal is known as prior, a confusion matrix could be produced to indicate the classification accuracy by our direct signal search approach. For comparison, we run BLAST (Altschul *et al.*, 1997) for each of the four 800 bp haplotype sequences against the read database, and collect those reads if the sequence identity and the length coverage is above 85% to all of the four sequences. For the resultant 40 reads, the haplotype is labeled based on the maximal BLAST bit score among the four haplotype sequences. As shown in Table 7, our signal-based algorithms achieved 100% accuracy, whereas the accuracy of read-based approach is lower than 90% (see Table 8). This result indicates that the haplotype classification at the raw-signal level is more accurate than that at the read level.

**Table 7.** The confusion matrix of haplotyping for MDM2 gene by signal-based approach

| Predicted \ Truth |  |  |  |  |
| --- | --- | --- | --- | --- |
|  | 285C-309T | 285C-309G | 285G-309T | 285G-309G |
| 285C-309T | 23 | 0 | 0 | 0 |
| 285C-309G | 0 | 0 | 0 | 0 |
| 285G-309T | 0 | 0 | 0 | 0 |
| 285G-309G | 0 | 0 | 0 | 21 |

**Table 8.** The confusion matrix of haplotyping for MDM2 gene by read-based approach

| Predicted \ Truth |  |  |  |  |
| --- | --- | --- | --- | --- |
|  | 285C-309T | 285C-309G | 285G-309T | 285G-309G |
| 285C-309T | 18 | 0 | 0 | 0 |
| 285C-309G | 1 | 0 | 0 | 1 |
| 285G-309T | 2 | 0 | 0 | 1 |
| 285G-309G | 0 | 0 | 0 | 17 |

## CONCLUSION AND DISCUSSION

We proposed two novel algorithms for local genome-to-signal search and mapping, which is a key step in major tasks of targeted sequencing. The proposed algorithms are based on the idea of subsequence dynamic time warping and directly operate on the nanopore raw signals. Comprehensive experiments on real-world datasets demonstrate that the proposed algorithms are able to produce accurate and efficient subsequence search, mapping and pattern classification. Two case studies further demonstrate the potential applications of our methods towards nanopore-based targeted sequencing.

Our proposed algorithms could also be extended and applied to detecting other single nucleotide variants (SNV), such as small insertions and deletions (InDels), as well as DNA modifications using nanopore data. Reports have shown that these events are challenging to detect, especially under a low sequencing coverage or with low-quality raw signals. As these non-standard events would all cause changes in the raw signals, it is possible to develop a universal detector for SNVs and DNA modifications under our framework. In addition, our algorithms are can be possibly used to resolve the long insertion and deletion events or other large scale mutation events with the help of large gap penalty (Smith and Waterman, 1981) or Viterbi-like algorithms (Viterbi, 2006) with hidden Markov models.

## ACKNOWLEDGEMENTS

The authors thank Minh Duc Cao, Lachlan J.M. Coin, Louise Roddam and Tania Duarte for providing the nanopore sequencing data. This work was supported by the King Abdullah University of Science and Technology (KAUST) Office of Sponsored Research (OSR) under Awards No. FCC/1/1976-04, URF/1/2601-01, URF/1/3007-01, URF/1/3412-01, URF/1/3450-01, URF/1/1976-26, and URF/1/1976-23.

