## Supplemental 1 for "Novel algorithms for efficient subsequence searching and mapping in nanopore raw signals towards targeted sequencing"

†These authors contributed equally.

### S1 The resampling technique based on FIR filter

As common knowledge, for the current MinION chemistry, ONT reports that single DNA strands are pulled through the pore at an average speed of 450 bp/s, while the electric current is sampled at a frequency of 4 kHz. This means that there are on average nine discrete measurements per $k$-mer (Rang *et al.*, 2018). Here, from the view of the signal process, the measured nanopore raw current signal is sampled in a redundant way. The encoding of a signal with redundant sampling could be reduced by the downsampling technique with little loss of information.

To preserve the information underlying the current signal while avoiding the aliasing, a resampling operation with a filter is adapted. Given a signal $X = (x_1, x_2, \cdots, x_N)$, the resampling operation on signal $X$ consists of the following three steps:

1. Upsample the signal $X$ by an integer factor $p$ (i.e., zero-insertion) to gain a signal $X_{up}$ with $p \cdot N$ time points.
2. Apply an FIR (finite-impulse response filter) to $X_{up}$ to get $X_{fir}$.
3. Downsample $X_{fir}$ (or decimate) by an integer factor $q$ (i.e., decimation), resulting in a signal $X_{down}$ with $p/q \cdot N$ time points.

Here, the FIR filter is a low pass filter designed with KaiserBessel window. An efficiently implemented polyphase filter bank based on the acceleration details in Saramaki and Bregovic (2002) is used to construct the low pass filter.

Because there are on average 9 discrete measurements per $k$-mer, here we use a $p = 1, q = 8$ setting to resample the raw current signal (thus, downsampling with 1/8 do not corrupt the Nyquist sampling theorem). Considering the high noise residing the nanopore raw current signal, the FIR resampling result $X_{down}$ will not be ideally accurate. However, in our signal detection and alignment task, a moderate level of resampling error is acceptable, which will be fixed by the subsequent multi-scale alignment schedule.

### S2 The existence of minimal length

**Lemma:** *A minimal length always exists for a set of nanopore raw signals that are collected in the same configuration.*

**Proof:** From the $k$-mer model provided by the Nanopore technology, a measured current signal $x$ belongs to one of the random variables $\{S_1, S_2, \cdots, S_N\}$, in which $S_i \sim \mathcal{N}(\mu_i, \sigma_i^2)$ for each $S_i$ ($N = 4^5$ for the 5-mer model and $4^6$ for the 6-mer model). For the convenience of further discussion, we assume that the variance of different $S_i$ has a similar value, i.e., $\forall i$, $\sigma \approx \sigma_i$.

If $x$ is a measured value of $S_i$ and $y$ is a measured value of $S_j$, the statistic of $x - y$ will belong to $D_{ij} = S_i - S_j$ and $D_{ij} \sim \mathcal{N}(\mu_i - \mu_j, \sigma_i^2 + \sigma_j^2)$, according to the property of Gaussian distribution. Specifically, we could find that $D_{ii} \sim \mathcal{N}(0, \sigma_i^2 + \sigma_j^2)$ if $i = j$.

Now consider the distance value of a DTW path $\{< i_1, j_1 >, < i_2, j_2 >, \cdots, < i_L, j_L >\}$ ($L$ is the length of the path; because omission is not considered in subsequence-extended DTW, here we assume that the length of detected paths for different locations are the same), in which

$$\text{Dist}(X, Y) = \sum_{k=1}^{L} |x_{i_k} - y_{j_k}|. \tag{1}$$

Here we could find that $\text{Dist}(X, Y)$ is also a random variables in statistic, which belongs to

$$C = \sum_{i=1}^{N} \sum_{j=1}^{N} (\alpha_{ij} |D_{ij}|), \tag{2}$$

where $\sum_{i=1}^{N}\sum_{j=1}^{N}\alpha_{ij} = L$ and $\alpha_{ij}$ is a positive integer value to indicate how many times $D_{ij}$ appears in the path. Because $D_{ij}$ follows a Gaussian distribution, the random variables $D_{ij}$ will have a folded normal distribution. For the convenience of further discussion, we define $\mu_{ij} = \mu_i - \mu_j$ and $\omega = 2\sigma^2 \approx \sigma_i^2 + \sigma_j^2$. Consequently,

$$\mu_{|D_{ij}|} = \omega\sqrt{\frac{2}{\pi}}\exp\left(\frac{-\mu_{ij}^2}{2\omega^2}\right) + \mu_{ij}\,\mathrm{erf}\left(\frac{\mu_{ij}}{\sqrt{2\omega^2}}\right), \tag{3}$$

and

$$\sigma_{|D_{ij}|}^2 = \mu_{ij}^2 + \omega^2 - \mu_{|D_{ij}|}^2, \tag{4}$$

where $\mathrm{erf}(z) = \frac{2}{\sqrt{\pi}}\int_0^z e^{-t^2}\mathrm{d}t$ is the error function. Obviously, $\mu_{|D_{ii}|} = \omega\sqrt{\frac{2}{\pi}}$ and $\sigma_{|D_{ii}|}^2 = (1-\frac{2}{\pi})\omega^2$ ($\mu_{ii} = 0$ for $i = j$). Because

$$\begin{aligned}\mu'_{|D_{ij}|}(\mu_{ij}) &= \omega\sqrt{\frac{2}{\pi}}\frac{d}{d\,\mu_{ij}}\exp\left(\frac{-\mu_{ij}^2}{2\omega^2}\right)\\ &+ \mathrm{erf}\left(\frac{\mu_{ij}}{\sqrt{2\omega^2}}\right)\frac{d}{d\,\mu_{ij}}\mu_{ij} + \mu_{ij}\frac{d}{d\,\mu_{ij}}\mathrm{erf}\left(\frac{\mu_{ij}}{\sqrt{2\omega^2}}\right)\\ &= \mathrm{erf}\left(\frac{\mu_{ij}}{\sqrt{2\omega^2}}\right),\end{aligned} \tag{5}$$

we can find that $\mu_{|D_{ij}|}$ decreases when $\mu_{ij} < 0$ and increases when $\mu_{ij} > 0$ (erf($\cdot$) is an odd function and $\geq 0$ in the right axis), which means that $\mu_{|D_{ij}|}$ gets the minimum value at $\mu_{ij} = 0$. Therefore, $\mu_{|D_{ii}|} \leq \mu_{|D_{ij}|}$ for all $i$ and $j$.

Now come back to the discussion of the DTW path. Here, we further introduce $\bar{\mu}_{|D_{ij}|} = \frac{1}{N^2}\sum_{i=1}^{N}\sum_{j=1}^{N}(\mu_{|D_{ij}|})$ and $\bar{\sigma}_{|D_{ij}|} = \frac{1}{N^2}\sum_{i=1}^{N}\sum_{j=1}^{N}(\sigma_{|D_{ij}|})$, and assume the elements in the mismatched paths are fully random (random variable $C_{mismatch}$) and the elements in the well-matched paths have $p$ sequence error ($0 \leq p \leq 1$) (random variable $C_{match}$). By relaxing the computation of exception $E(C)$, we can get that

$$E(C_{mismatch}) = L\bar{\mu}_{|D_{ij}|}, \tag{6}$$

and

$$E(C_{match}) = p \cdot L\bar{\mu}_{|D_{ij}|} + (1-p)L\mu_{|D_{ii}|}. \tag{7}$$

Obviously, $E(C_{mismatch}) - E(C_{match}) = (1-p)L(\bar{\mu}_{|D_{ij}|} - \mu_{|D_{ii}|})$. The safe value of $L$ can be constructed as

$$L = \frac{Q}{(1-p)(\bar{\mu}_{|D_{ij}|} - \mu_{|D_{ii}|})} \tag{8}$$

to make short reads be detected with high confidence ($Q$ is a parameter to indicate the sensitivity, i.e., at what value a peak should be judged distinguishable). Here, the value of $L$ is the *minimal length* we need.

### S3 Experimental details for the SNP detection

Here, a case study is shown to demonstrate how we can identify and visualize SNPs based on the nanopore raw signal data produced by the similarity searching and exact mapping of our algorithm. The experiment is carried out on the E. coli dataset with a relatively low coverage (20×).

Above all, we first generate the synthetic reference genomes by randomly substituting 100 bases on the sense strand of the E. coli reference genome, as shown in Figure S1.

Then we extract a 2000-long sequence on the synthetic genome that might contain a SNP and search this reference sequence as a query against the nanopore signal database by using our proposed algorithm cwSDTWnano, to localize the candidate raw signal segments and positions that might contain a SNP, as shown in Figure S2.

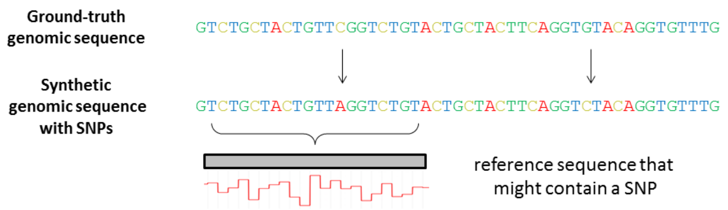


Figure S1: Experimental design for the single nucleotide polymorphisms (SNP) detection. A synthetic reference E. coli genome is generated by randomly substituting K bases on the sense strand from the ground-truth genome. Then this synthetic genomic sequence serves as our reference, and we may find evidence for a subsequence that might contain a SNP from the nanopore signal database.

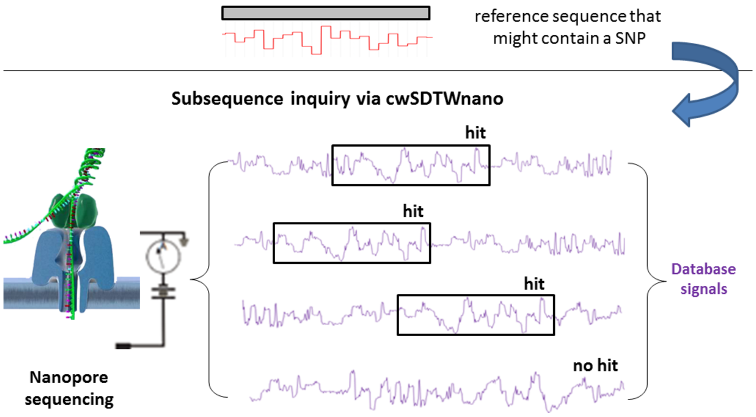


Figure S2: Subsequence inquery via cwSDTWnano. For a query reference sequence that might contain a SNP, we first transfer the query sequence to the expected signal by the k-mer pore model, and search this query signal against the nanopore signal database via cwSDTWnano. Note that we filter out those low quality database signals whose normalized distance to the query signal is large. These 'hit' signals might serve as the evidence to determine the SNP location as well as the SNP type.

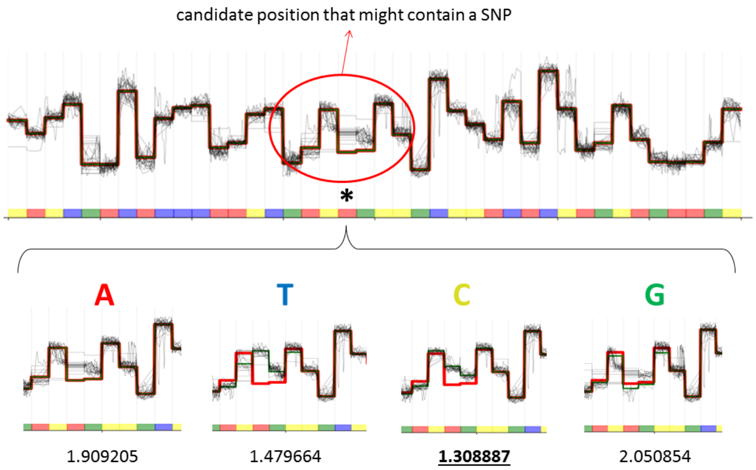


Figure S3: Procedure to locate candidate SNP positions and the SNP type. The candidate positions that might contain a SNP are chosen based on the mismatches between the aligned 'hit' signals and the query reference sequence's $k$-mer pore model. Afterward, for each candidate position, four genomic sequences centered at this position with four possible nucleotides A,T,C,G are selected as the query sequences to search the signal database again. Finally, the sequence containing the nucleotide whose expected signal (i.e., $k$-mer pore model) is closest to the observed signals in the database is chosen as the possible SNP type at the candidate position. In particular, the expected signal derived from the reference sequence is shown in the bold red curve, whereas those expected signals derived from the other three sequences are shown in the thin green curve. It is obvious that the expected signal from a sequence containing nucleotide is the most similar to the observed signals, which will be the SNP type at this position.

These candidate positions are chosen based on the mismatches between the aligned signals and the reference sequence's $k$-mer pore model. Afterward, for each candidate position, four genomic sequences with length 600 centered at this position with four possible nucleotides $\{A, T, C, G\}$ are selected as the query sequences to search against the signal database again. Note that we filter out those database signals whose normalized distance is larger than a given threshold (by default is 0.20) to the expected signal that is translated from the query sequence (based on the 6-mer pore model). Finally, the mutation (nucleotide $X, X \in \{A, T, C, G\}$) with the expected signal closest to the observed signals in the database are chosen as the possible SNP at the candidate position, as shown in Figure S3.

A representative example of SNP identification is visualized in Figure S4, which is the mapping of a set of aligned raw signals. Here the red (green) curve indicates the 6-mer pore model for the reference (mutated) sequence centered at the candidate SNP position.


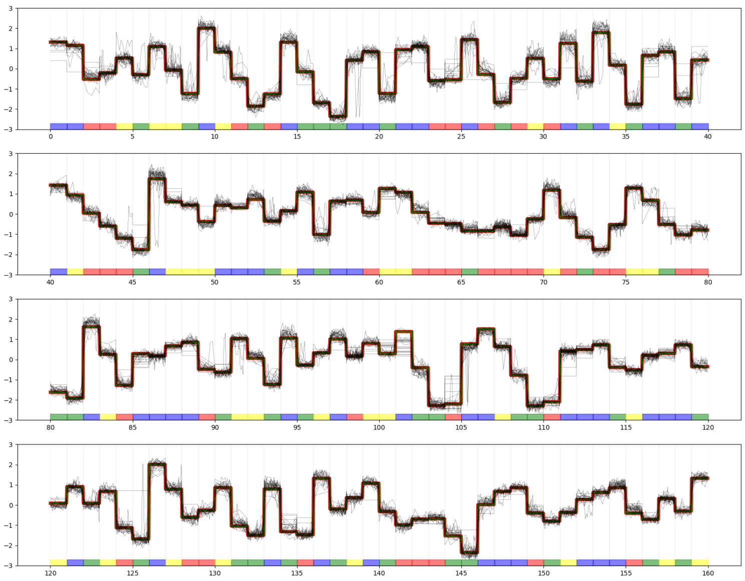

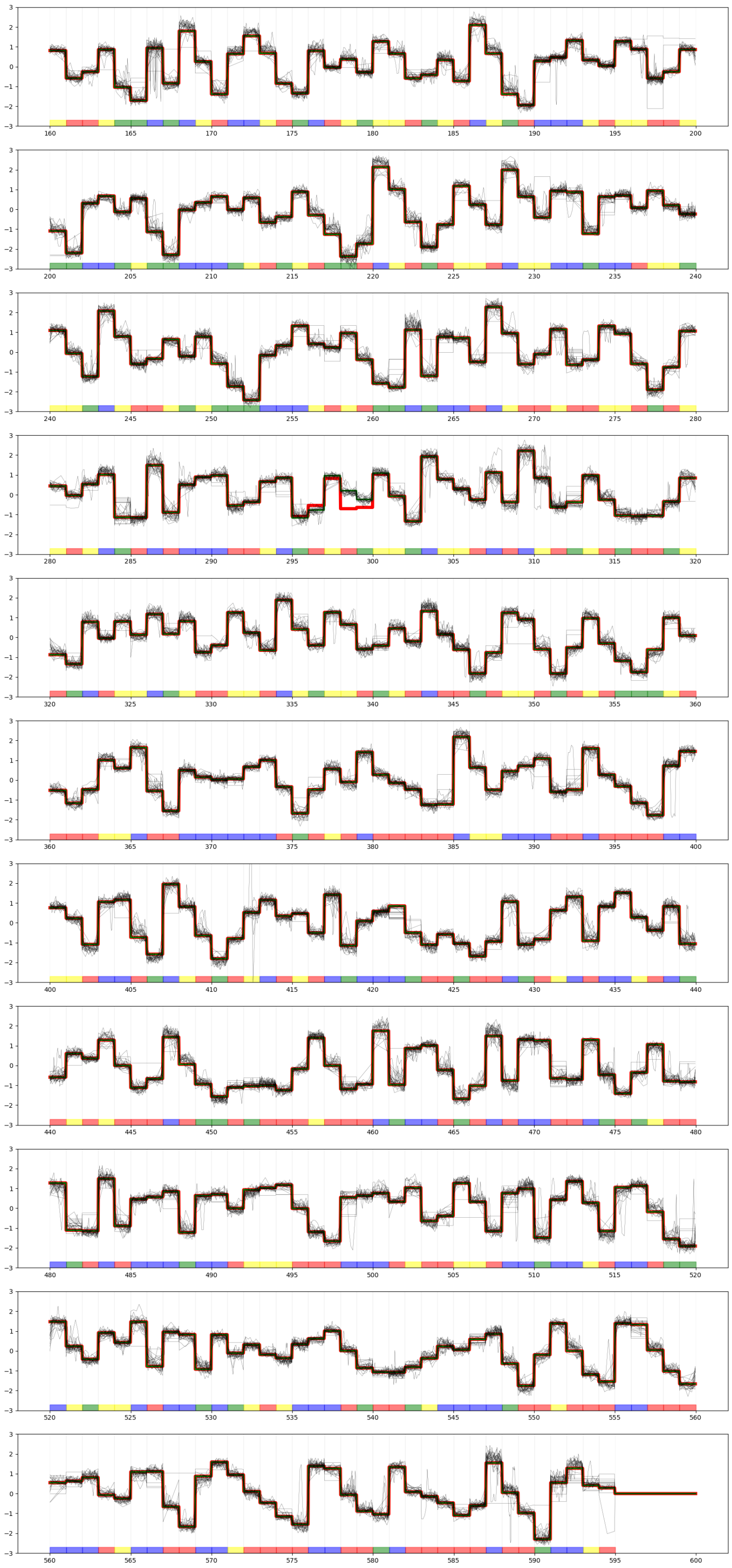

Figure S4: The visualization of the identified SNP (locate at [295:300]) by cwSDTWnano, which is a segmentation from a set of aligned raw signals. Here, the nucleotide A, C, G and T on the reference sequence (query) are labeled by red, yellow, green and blue color, respectively. The aligned nanopore signals are shown in black squiggle curves and the red (green) curve indicates the 6-mer pore model for the reference (mutated) sequence centered at the candidate of SNP position.
